# Interaction of intraocular pressure and ganglion cell function in open angle glaucoma

**DOI:** 10.1101/2020.01.30.924290

**Authors:** K. O. Al-Nosairy, J.J.O.N. Van den Bosch, V. Pennisi, H. Thieme, K Mansouri, L. Choritz, M. B. Hoffmann

## Abstract

**Purpose:** To test the feasibility of simultaneous steady-state pattern electroretinogram (PERG) and intraocular pressure (IOP) measurements with an IOP sensor and to test a model for IOP manipulation during lateral decubitus positioning (LDP) and its impact on the PERG.

**Design:** A prospective, observational study.

**Methods:** 15 healthy controls and 15 treated glaucoma patients participated in the study. 8 patients had an intraocular IOP sensor (eyemate-IO^®^, Implandata Ophthalmic Products GmbH) in the right eye (GLA_IMP_) and 7 had no sensor and with glaucoma in the left eye. (1) We tested the feasibility of simultaneous IOP and PERG recordings by comparing PERGs with and without simultaneous IOP-read out in GLA_IMP_. (2) All participants were positioned in the following order: sitting1 (S1), right LDP (LD_R_), sitting2 (S2), left LDP (LD_L_) and sitting3 (S3). For each position, PERG amplitudes and IOP were determined with rebound tonometry (Icare® TA01i) in all participants without the IOP sensor.

**Results:** Electromagnetic intrusions of IOP sensor readout onto steady-state PERG-recordings had, due to different frequency ranges, no relevant effect on PERG amplitudes. IOP and PERG measures were affected by LDP, e.g., IOP was increased during LD_R_ vs S1 in the lower eyes of GLA_IMP_ and controls (P < 0.001 and P < 0.05, respectively) and PERG amplitude was decreased (P < 0.05 and P < 0.01, respectively).

**Conclusions:** During LDP, IOP and PERG measurements changed more in the lower eye. IOP changes induced by LDP may be a model for studying the interaction of IOP and ganglion cell function.

## Introduction

While glaucoma is a global leading cause of irreversible blindness^1–3^, its pathogenesis is not yet fully understood. One approach to fill this gap is deciphering the mechanisms via which intraocular pressure (IOP), a well-known risk for glaucoma, impacts on ganglion cell function. An attractive maneuver to elucidate these mechanisms is the manipulation of IOP, e.g. by utilizing posture-induced IOP-changes. And to manifest its impact on retinal function, e.g. with non-invasive electrophysiology.

An important tool for the assessment of retinal ganglion cell function is the steady state pattern electroretinogram (ssPERG)^4,5^. Consequently, the PERG is of value for the detection and investigation of glaucoma^6,7^. Previous studies indicate that PERG-measurements during IOP manipulations may be of diagnostic importance in glaucoma and have a predictive role for future conversion of glaucoma suspects. Specifically, posture-induced IOP changes offer a straightforward option to manipulate IOP. For example, supine posture with −10° head down tilt in a subpopulation of glaucoma suspects was found to be associated with an increase in IOP and a decrease of PERG amplitude^8,9^. An increase in IOP was also reported during lateral decubitus posture (LDP) for both participants with healthy vision^10–12^ and for glaucoma patients^13–15^. Such approaches would greatly benefit from simultaneous IOP and PERG measurements, which was up-to-date technically not feasible. Recently developed telemetric IOP monitoring devices have the potential to fill this gap, as they open the possibility of simultaneous IOP- and electrophysiological recordings.

Telemetric IOP sensors were first proposed in 1967^16^ but were not tested in clinical trials. At present, two technologies are commercially available for continual IOP monitoring: a contact lens sensor (Sensimed Triggerfish)^17–19^ and an implantable intraocular pressure sensor (eyemate-IO sensor)^20–23^. With the latter approach, a wireless ring shaped sensor is placed in the ciliary sulcus during co-implantation of an intraocular lens during cataract surgery^21^. Studies have shown the eyemate-IO sensor to be safe, well tolerated and of good functionality to continuously measure IOP^21,23–25^. In principle, it should enable simultaneous PERG and IOP readings.

This study is the first to perform simultaneous electroretinographical recordings and continual IOP measurements in glaucoma patients. We assessed the feasibility of simultaneous PERG and IOP measurements and demonstrated the relation of ganglion cell responses to IOP changes in LDP, with the lower eye, i.e also termed ‘dependent eye’^12^), being most affected during LDP.

## Methods

### Participants

This prospective observational study was conducted in University Eye Clinic of Otto-von-Guericke University of Magdeburg. The participants gave their written consent to participate in the study. The procedures followed the tenets of the declaration of Helsinki and the protocol was approved by the ethical committee of the Otto-von-Guericke University of Magdeburg, Germany. All participants, three groups as detailed below, underwent complete ophthalmic examinations and best corrected visual acuity testing using the Early Treatment for Diabetic Retinopathy Study’s (ETDRS) chart^26^ and visual field testing using the Swedish Interactive Threshold Algorithm 24-2 protocol (SITA) of Humphrey Field Analyzer 3 (Carl Zeiss Meditec AG, Jena, Germany). Clinical and demographic data for all participants are shown in Table 1.

**Table 1.**
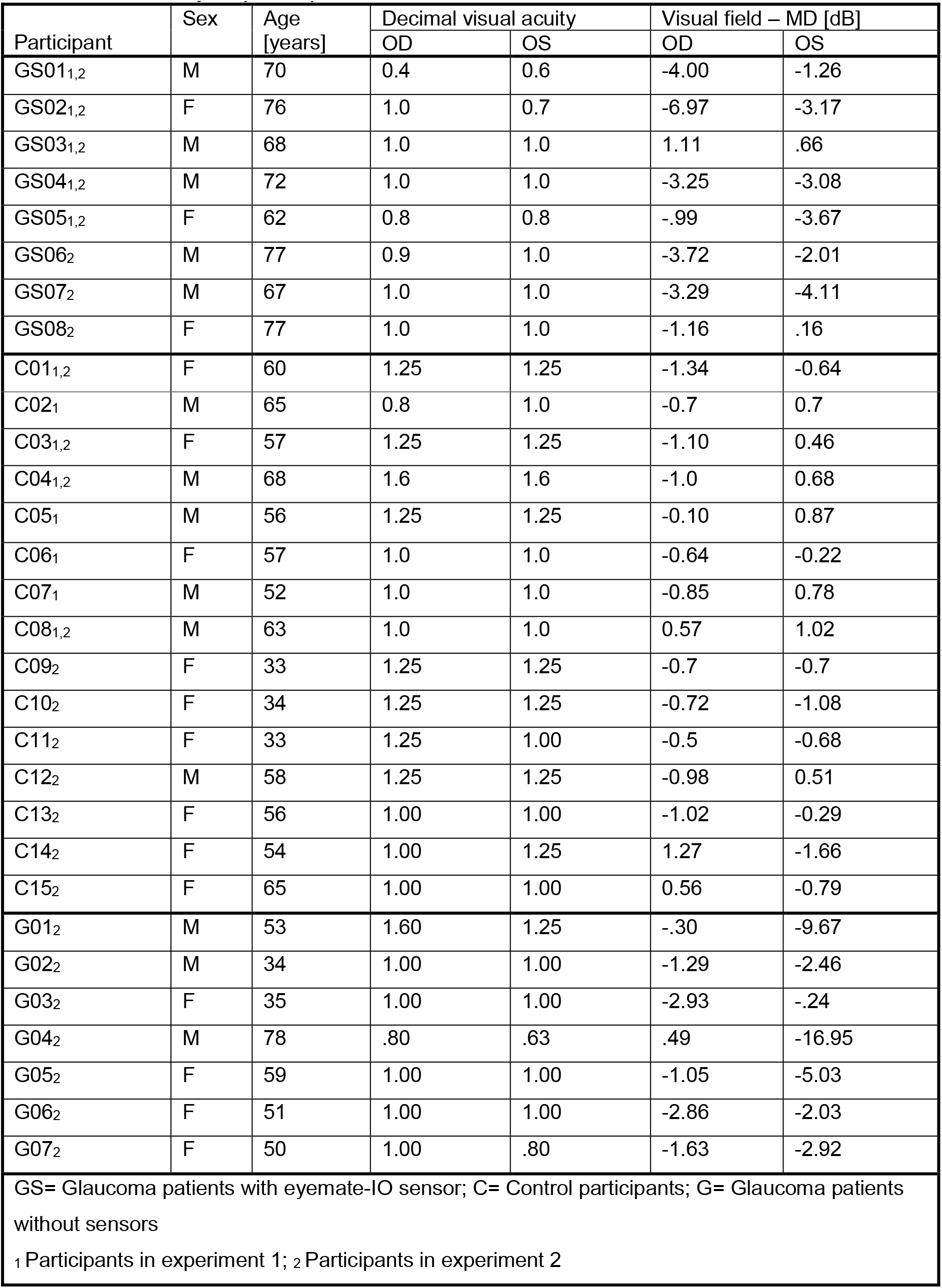
Summary of participant characteristics.

| Participant | Sex | Age<br>[years] | Decimal visual acuity |  | Visual field – MD [dB] |  |
| --- | --- | --- | --- | --- | --- | --- |
|  |  |  | OD | OS | OD | OS |
| GS01<sub>1,2</sub> | M | 70 | 0.4 | 0.6 | -4.00 | -1.26 |
| GS02<sub>1,2</sub> | F | 76 | 1.0 | 0.7 | -6.97 | -3.17 |
| GS03<sub>1,2</sub> | M | 68 | 1.0 | 1.0 | 1.11 | .66 |
| GS04<sub>1,2</sub> | M | 72 | 1.0 | 1.0 | -3.25 | -3.08 |
| GS05<sub>1,2</sub> | F | 62 | 0.8 | 0.8 | -.99 | -3.67 |
| GS06<sub>2</sub> | M | 77 | 0.9 | 1.0 | -3.72 | -2.01 |
| GS07<sub>2</sub> | M | 67 | 1.0 | 1.0 | -3.29 | -4.11 |
| GS08<sub>2</sub> | F | 77 | 1.0 | 1.0 | -1.16 | .16 |
| C01<sub>1,2</sub> | F | 60 | 1.25 | 1.25 | -1.34 | -0.64 |
| C02<sub>1</sub> | M | 65 | 0.8 | 1.0 | -0.7 | 0.7 |
| C03<sub>1,2</sub> | F | 57 | 1.25 | 1.25 | -1.10 | 0.46 |
| C04<sub>1,2</sub> | M | 68 | 1.6 | 1.6 | -1.0 | 0.68 |
| C05<sub>1</sub> | M | 56 | 1.25 | 1.25 | -0.10 | 0.87 |
| C06<sub>1</sub> | F | 57 | 1.0 | 1.0 | -0.64 | -0.22 |
| C07<sub>1</sub> | M | 52 | 1.0 | 1.0 | -0.85 | 0.78 |
| C08<sub>1,2</sub> | M | 63 | 1.0 | 1.0 | 0.57 | 1.02 |
| C09<sub>2</sub> | F | 33 | 1.25 | 1.25 | -0.7 | -0.7 |
| C10<sub>2</sub> | F | 34 | 1.25 | 1.25 | -0.72 | -1.08 |
| C11<sub>2</sub> | F | 33 | 1.25 | 1.00 | -0.5 | -0.68 |
| C12<sub>2</sub> | M | 58 | 1.25 | 1.25 | -0.98 | 0.51 |
| C13<sub>2</sub> | F | 56 | 1.00 | 1.00 | -1.02 | -0.29 |
| C14<sub>2</sub> | F | 54 | 1.00 | 1.25 | 1.27 | -1.66 |
| C15<sub>2</sub> | F | 65 | 1.00 | 1.00 | 0.56 | -0.79 |
| G01<sub>2</sub> | M | 53 | 1.60 | 1.25 | -.30 | -9.67 |
| G02<sub>2</sub> | M | 34 | 1.00 | 1.00 | -1.29 | -2.46 |
| G03<sub>2</sub> | F | 35 | 1.00 | 1.00 | -2.93 | -.24 |
| G04<sub>2</sub> | M | 78 | .80 | .63 | .49 | -16.95 |
| G05<sub>2</sub> | F | 59 | 1.00 | 1.00 | -1.05 | -5.03 |
| G06<sub>2</sub> | F | 51 | 1.00 | 1.00 | -2.86 | -2.03 |
| G07<sub>2</sub> | F | 50 | 1.00 | .80 | -1.63 | -2.92 |
| GS= Glaucoma patients with eyemate-IO sensor; C= Control participants; G= Glaucoma patients without sensors |  |  |  |  |  |  |
| <sub>1</sub> Participants in experiment 1; <sub>2</sub> Participants in experiment 2 |  |  |  |  |  |  |

Exclusion criteria were any systemic diseases, ocular diseases or surgeries that might affect electrophysiology recordings except cataract surgery and, in the glaucoma group, glaucoma surgery or incipient cataract that did not decrease BCVA < 0.8^27^ and refractive error exceeding −6 or +3 D or astigmatism ≥ 2 D.

#### GLA_IMP_ group

Eight patients (age range [years]: 62-77 years; mean ± SE: 71.1 ± 1.9 years) with open angle glaucoma who had previously been implanted with a telemetric IOP sensor in the right eye (eyemate-IO^®^, Implandata Ophthalmic Products GmbH, Hannover, Germany), were enrolled in the study. Five of 8 subjects participated in experiment 1 to test the influence of eyemate-IO sensor electromagnetic radiation on PERG recordings in both eyes. All 8 participants participated in experiment 2 to determine the LDP effect on PERG amplitudes. Intraocular pressure and PERG measurements were included only for the right eye as not all left eyes had glaucoma. The eyemate-IO sensor had previously (at least 3 years prior to the present measurements) been implanted in glaucomatous patients as part of a preceding study^24^.

#### GLA_LE_-group

The left eyes of 7 patients (age range [years]: 34-78; mean age ± SE: 51.4 ± 5.6) with open angle glaucoma (POAG) were included and were compared to controls’ left eyes (CON_LE_ as defined below). Not all right eyes of this group had glaucoma; right eyes were, therefore, not included in the analysis.

All OAG eyes either had glaucoma hemifield test that was outside normal limits (Humphrey Field Analyzer 3, as detailed below), a typical visual field glaucomatous damage manifested as a cluster of 3 or more non-edge points all depressed on the pattern deviation plot < 5% and one of which depressed < 1% or abnormal corrected pattern standard deviation < 5% on Humphrey Swedish interactive threshold algorithm 24-2 (SITA fast)^28^ and a glaucomatous appearance of the optic disc, i.e. with a general enlargement of the cupping defined as vertical cup-to-disc ratio ≥ 0.7, retinal fiber layer defect or a local notching of the rim and an open anterior chamber angle. All glaucoma patients were under IOP-lowering treatment.

#### Normal controls (CON_RE_/CON_LE_)

Fifteen subjects (age range [years]: 33-65; mean age ± SE: 52.4 ± 3.8) with normal visual acuity (see Table 1) without ocular diseases were included in the study. Eight were included in experiment 1; eleven were enrolled in experiment 2. In experiment 2, controls’ right eyes (CON_RE_) were compared to the right eye of GLA_IMP_ group whereas left eyes of controls (CON_LE_) were compared to GLA_LE_ group.

### IOP sensor (eyemate-IO sensor) and external reader device (Mesograph)

The eyemate-IO sensor is a <u>M</u>icro-<u>E</u>lectro-<u>M</u>echanical <u>S</u>ystem <u>A</u>pplication-<u>S</u>pecific <u>I</u>ntegrated <u>C</u>ircuit (MEMS ASIC) comprising pressure sensitive sensor cells, temperature sensors, analog-to-digital converters, and telemetry. The ASIC is bonded to a gold-made circular micro-coil antenna, both parts are silicon-encapsulated^29^. For readout of the sensor, typically an external reader device (Mesograph) is held in front of the eye with < 5 cm distance to power the implant. Upon button-press on the device, it emits a radio frequency field at 13.56 MHz, which establishes an electromagnetic link towards the implant. When sufficient power is available to the ASIC, a pressure reading is being performed and digitized data is transferred back to the external reader device. The eyemate-IO sensor can obtain IOP measurements comprising around 10 individual samples per second^21,29^. As an alternative to holding the external reader device manually in front of the eye, a reader can be connected to a coiled circular antenna that is positioned and attached around the eye with a plaster (Figure 1). This way, a continual IOP readout can be performed^30^ at an average sampling rate of 9.2 Hz and recorded with a computer via a USB connection. We applied this continuous readout mode in our study.

**Figure 1.**
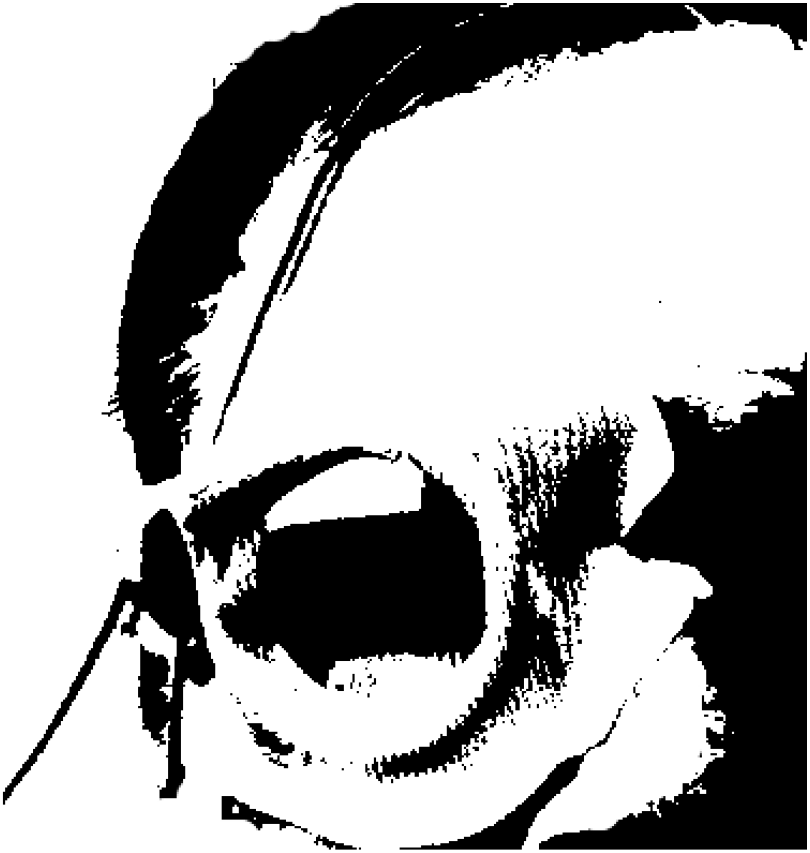
Recording setup for a GLA_IMP_ patient with binocular electrodes for PERG recordings and the antenna around the right eye for simultaneous continuous read-out of IOP responses in Experiments 1 & 2. (Permission obtained from patient)

### Visual stimuli, procedure and recordings

The EP2000 evoked potential system was used for stimulation, recording and analysis of steady-state PERGs^31^ following the international society for clinical electrophysiology of vision (ISCEV) standards for PERG-recordings^32^. The stimuli were presented binocularly at a frame rate of 75 Hz on a monochrome monitor (MDG403, Philips; P45 phosphor) in a dimly-lit room. The participants maintained fixation at the center of the monitor using a fixation cross (1° diameter) which was replaced by a 200 ms-duration digit randomly appearing every 5-20 seconds. Contrast-inverting (15 Hz) checkerboard patterns (visual field: 25° × 25°; mean luminance: 45 cd/m^2^; contrast: 98%) with two check sizes, 0.8° and 15°, were presented for stimulation at a viewing distance of 57 cm. Two PERG blocks were recorded for each sitting position and 4 PERG blocks were recorded for each LDP. Repeated blocks were averaged offline using Igor (IGOR Pro, WaveMetrics, Portland). Each PERG block contained 8 stimulus cycles of 10 trials per checksize (80 sweeps of 1.066 s trail duration). Signals exceeding ±90 μV were considered artifacts originating from eye movements or blinking, rejected and recollected. The pupils were not dilated.

Five silver-cup skin electrodes (9 mm diameter; Silver EEG Cup Electrodes, Natus Manufacturing Limited, Ireland) filled with conductive paste (Ten20, WEAVER and Company, USA) were used as recording and reference electrodes and applied after cleaning the skin with a cleaning paste (skinPure, NIHON KODEN Corporation, Tokyo, Japan) to keep skin conductance below 5 kOhm. Binocular PERGs were recorded using active skin electrodes placed about 5 mm below the eyelids vertically aligned with the pupil in the primary gaze^6,33^. The reference electrode was attached to the temple ipsilateral to the corresponding eye. Similarly, the ground electrode was placed at Fpz^34,35^. The signal was amplified by 50 k (Grass Model 12, Astro-Med, Inc., West Warwick, RI, USA), band-pass filtered 1-100 Hz and digitized at 1 kHz with 12-bit resolution by a G4 Power Macintosh computer running the EP2000 Evoked Potentials System^31^.

### Experiment 1 – Effect of eyemate-IO sensor read-out on PERG recordings

Since the eyemate IO-sensor functions employ electromagnetic radiation coupling with the antenna, we tested for possible interferences and effects of eyemate-IO sensor on the PERG recordings. To confirm any changes detected in patients, we additionally simulated IOP readout in controls, which required, due to the read-out procedure, the reader to detect a sensor, which was therefore placed externally next to the eye, i.e. without IOP-functionality. For this purpose, the eyemate-IO sensor was fixed below the right lateral third of the lower eyelid with the antenna placed in front of it to allow electromagnetic coupling to the external reader. Participants in experiment 1 underwent four blocks of PERG binocular recordings, two with read-out switch on, i.e. IO-Reader_ON_, and two with read-out switched off, i.e. IO-Reader_OFF_, in a counterbalanced sequence (‘A-B-B-A’-scheme). The two PERG blocks (2 × 80 sweeps) per condition and same readout status were averaged. To assess the raw electrophysiological recordings prior to averaging in EP2000, the recordings were acquired using the PowerLab recording system (Model M880, ADInstrument Pty Ltd, Australia).

### Experiment 2 – Effect of lateral decubitus posture (LDP) on PERG

PERGs were recorded binocularly while the participants were positioned in the following sequence: Sitting (S1), right LDP (LD_R_), sitting (S2), left LDP (LD_L_) and sitting (S3). Five minutes after taking each position, the IOP was measured with an Icare tonometer (Icare® TA01i Tonometer, Helsinki, Finland) for the control- and the GLA_LE_-group or simultaneously during PERG recording with eyemate-IO sensor for the GLA_IMP_-group. Pillows were used to support the head during lateral decubitus and to assure that the head was parallel to the ground. Care was taken that the pillow did not compress the lower eye during LDP. The IOP was measured in the center of the cornea and started always with the right eye followed by the left eye; the IOP value was an automatic average of one set comprising 6 measurements. As a measure to reduce the probability of IOP erroneous readings (i.e tilting of the device and misalignment), we asked the patient to always look straight and kept the orientation of Icare TA01i parallel to the ground in both sitting position and LDP. Also, the use of Icare TA01i is clinically robust and appeared insensitive to lateral and angular deviations during measurements^36^.

### Analysis and statistics

ssPERG were Fourier-analyzed and the response amplitude at stimulation frequency, i.e. 15 Hz was determined and corrected for the noise estimate, i.e. an average of the neighboring frequencies below and above 15 Hz^37,38^.

The non-averaged traces acquired during experiment 1 were digitized with a PowerLab recording system (Model M880, ADInstrument Pty Ltd, Australia) and exported to Igor (IGOR Pro, WaveMetrics, Portland) for subsequent analyses and the determination of intrusion frequency during IOP readout from the eyemate-IO sensor. ssPERG amplitudes were transferred into IGOR sheet containing glaucoma and controls data and graphic representation of each amplitude check size in both eyes and both reader-out states were plotted. Similarly, ssPERG amplitudes were plotted in Igor for controls and glaucoma_IMP_ patients for different postures. To test the effect of LDP on IOP and ganglion cell function, these measures were compared for S1 vs LD_R_ and S2 vs LD_L_ with paired T-tests using SPSS 24 (statistical Package for the Social Sciences, IBM). P values were corrected using the Bonferroni-Holm correction^39^ for multiple comparisons.

## Results

### Experiment 1 – Feasibility of simultaneous eyemate-IO sensor read-out and ssPERG recordings

A qualitative overview over the ssPERG recordings with (IO-Reader_ON_) and without simultaneous eyemate-IO sensor read-out (IO-Reader_OFF_) in given in Figure 2 for a participant with the eyemate-IO sensor implanted. In Figure 2A non-averaged recording traces for IO-Reader_ON_ and IO-Reader_OFF_ are depicted. It is evident that there is a periodic high amplitude intrusion into the recordings only for IO-Reader_ON_, which was in the order of 80 μV (peak-to-peak) and at a frequency centred around 9.2 Hz, i.e. the known approximate readout frequency (see Methods). A set of averaged ssPERGs from the same participant is given in Figure 2B for both check sizes, for both conditions (IO-Reader_ON_ and IO-Reader_OFF_), and both eyes (with and without sensor implant for the right and left eye, respectively). Significant responses (P < 0.01) are evident at the stimulation frequency (15Hz) for all ssPERGs obtained, as underlined by the peak in the spectrum at 15 Hz. For the IO-Reader_ON_ condition, a second peak is observed at around 9 Hz, i.e. corresponding to the read-out frequency (see Methods and Results above). It is also evident that there is an overspill of this intrusion to neighboring frequencies^37^. This can be attributed to slight variations in the read-out frequency and in an ssPERG averaging that is not locked to the read-out. These intrusions are expected to have minimal effect on the amplitude obtained at the stimulation frequency of 15 Hz, as it is not a directly neighboring frequency, and as noise estimates allow for the subtraction of the noise from the actual response amplitude^38^.

**Figure 2.**
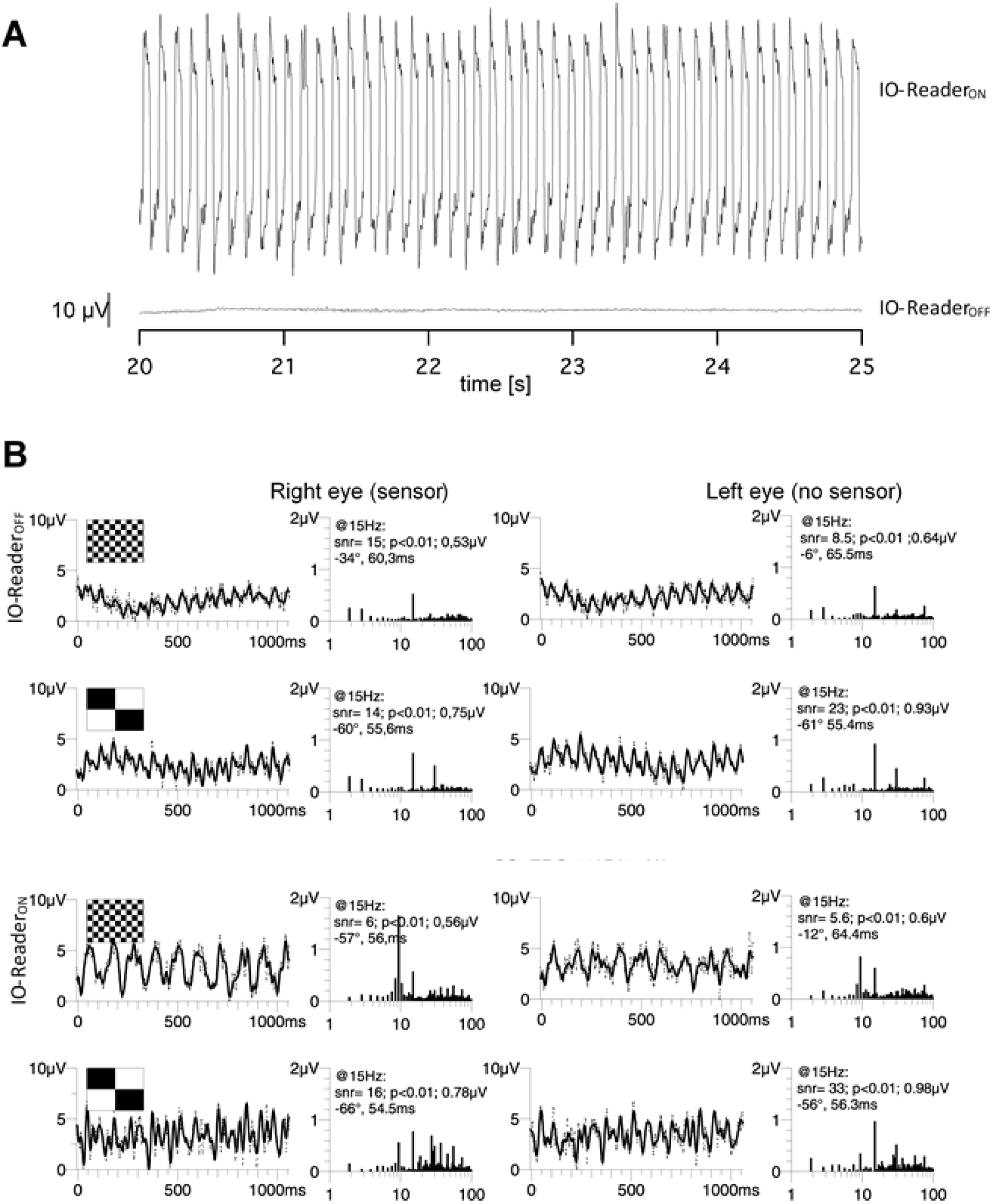
Effect of IO-readout on PERG recordings. (A) Raw PERG recording trace for eyemate-IO sensor on/off states. For IO-Reader_ON_ (top trace) the applied electromagnetic field induces a noise intrusion of around 9 Hz, i.e. at the read-out frequency (see Methods). For IO-Reader_OFF_ (bottom trace) the noise intrusion is absent. (B) ssPERGs and frequency spectra for IO-Reader_ON/OFF_ -states for both eyes for two check sizes (0.8° and 15°). Noise intrusions are reduced due to non-phase locked averaging, but still evident for both the eye with eyemate-IO sensor and - Reader and for the fellow eye. As a consequence, a response at around 9.2 Hz is evident in the frequency spectra for IO-Reader_ON_, in addition to the stimulus evoked response at 15 Hz.

To test this directly, we performed a quantitative comparison of ssPERG-amplitudes for IO-Reader_ON/OFF_ in a total of 5 participants with an eyemate-IO implant and in an additional 8 controls without the implant, but use of the IO-Reader (see Methods). For this purpose we analyzed the effect of the conditions IO-Reader_ON/OFF_ on ssPERG-amplitudes for both check sizes, i.e. 0.8° and 15° as depicted in Figure 3. On average only small trends were observed, which did not reach significance with T-test statistical testing for neither eye nor checksizes as shown in Table 2 for the right eyes of GLA_IMP_ group and of controls with attached eyemate-IO sensor/reader antenna. Taken together, these results indicate a lack of relevant impact of the eyemate-IO sensor on ssPERG recordings.

**Table 2.**
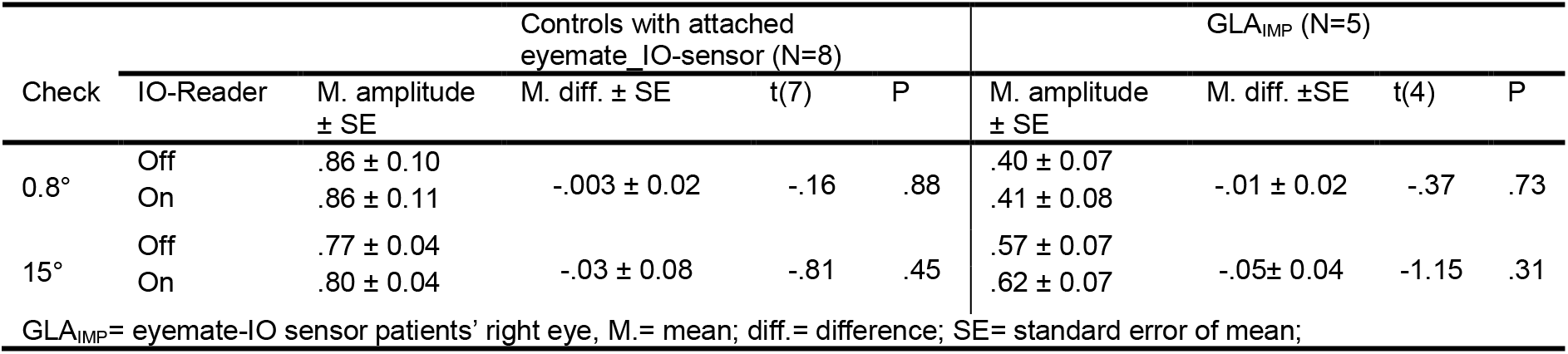
PERG amplitudes comparisons between the IO-Reader_Off/ON_ states in right eye of controls with attached eyemate-IO sensor and right eye of glaucoma patients with eyemate-IO implant.

**Figure 3.**
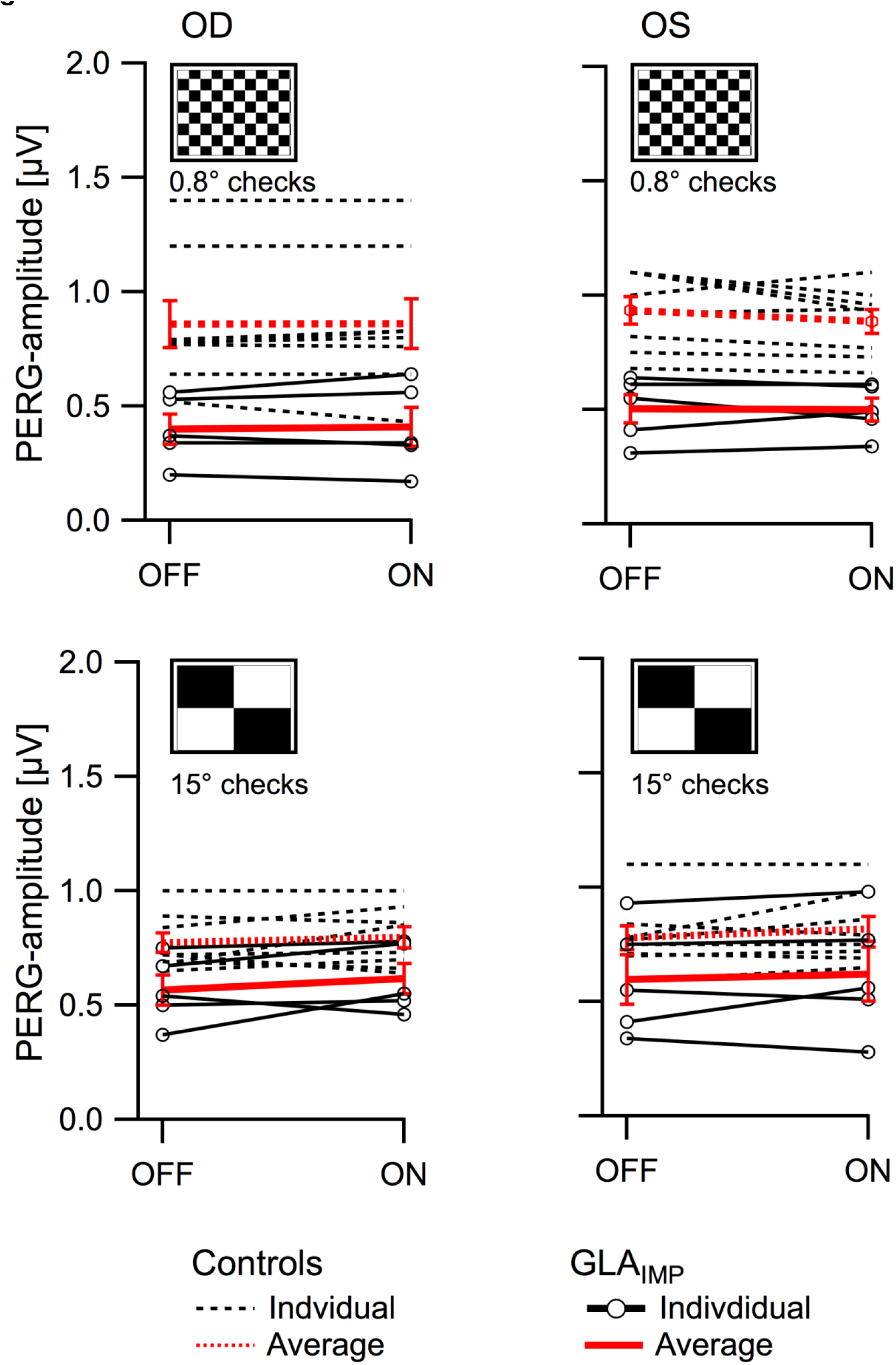
PERG amplitudes for 0.8° and 15° checksizes and IO-Reader_ON_ and IO-Reader_OFF_ in GLA_IMP_ (n=5) and controls (n=8). No significant differences between IO-Reader_ON_ and IO-Reader_OFF_ were evident. Average data ±SEM (red) and single subject effects are depicted. GLA_IMP_: Glaucoma patients with eyemate-IO implant.

### Experiment 2 – Influence of postural change on IOP

We investigated the effect of posture (lateral decubitus vs. sitting) on IOP and PERG amplitudes as depicted in Figure 4. We were particularly interested in these effects in the glaucomatous eyes of the participants with simultaneous IOP and PERG-readout, i.e. the right eye of the GLA_IMP_ group (Figure 4 A). For GLA_IMP_, the IOP increased for LD_R_ vs S1 by 5.1 mmHg (P < 0.001), and for LD_L_ vs S2 by 2.7 mmHg (P = 0.01; Table 3). In CON_RE_, the IOP increased by 1.5 mmHg for LD_R_ vs S1 (P < 0.05) and for LD_L_ vs S2 (P < 0.05). In summary, effects of posture on IOP were evident for both groups and particularly pronounced for LD_R_ in GLA_IMP_ group, i.e. IOP increase in the lower eye.

**Table 3.** Intraocular pressure of right eye during different body postures

| Posture | CON<sub>RE</sub> (N=11) |  |  |  | GLA<sub>IMP</sub> (N=8) |  |  |  |
| --- | --- | --- | --- | --- | --- | --- | --- | --- |
|  | Mean IOP ± SE | M. diff. ± SE | t(10) | P | Mean IOP ± SE | M. diff. ± SE | t(7) | P |
| S1 | 15.36 ± 1.04 |  |  |  | 15.98 ± 1.79 |  |  |  |
| LD<sub>R</sub> | 16.91 ± 0.89 | -1.55 ± 0.58 | -2.68 | 0.023 | 21.12 ± 1.57 | -5.14 ± 0.57 | -8.96 | < .001 |
| S2 | 14.00 ± 1.10 |  |  |  | 16.97 ± 1.72 |  |  |  |
| LD<sub>L</sub> | 15.45 ± 1.23 | -1.45 ± 0.62 | -2.33 | 0.04 | 19.68 ± 1.91 | -2.71 ± 0.78 | -3.47 | 0.01 |

CON<sub>RE</sub>= Controls' right eye, GLA<sub>IMP</sub>= eyemate-IO sensor patients' right eye, M. diff.= mean difference; SE= standard error of mean; IOP= intraocular pressure; S1& S2= sitting 1& 2; LD<sub>R</sub> = right lateral decubitus posture; LD<sub>L</sub> = left lateral decubitus posture

**Figure 4.**
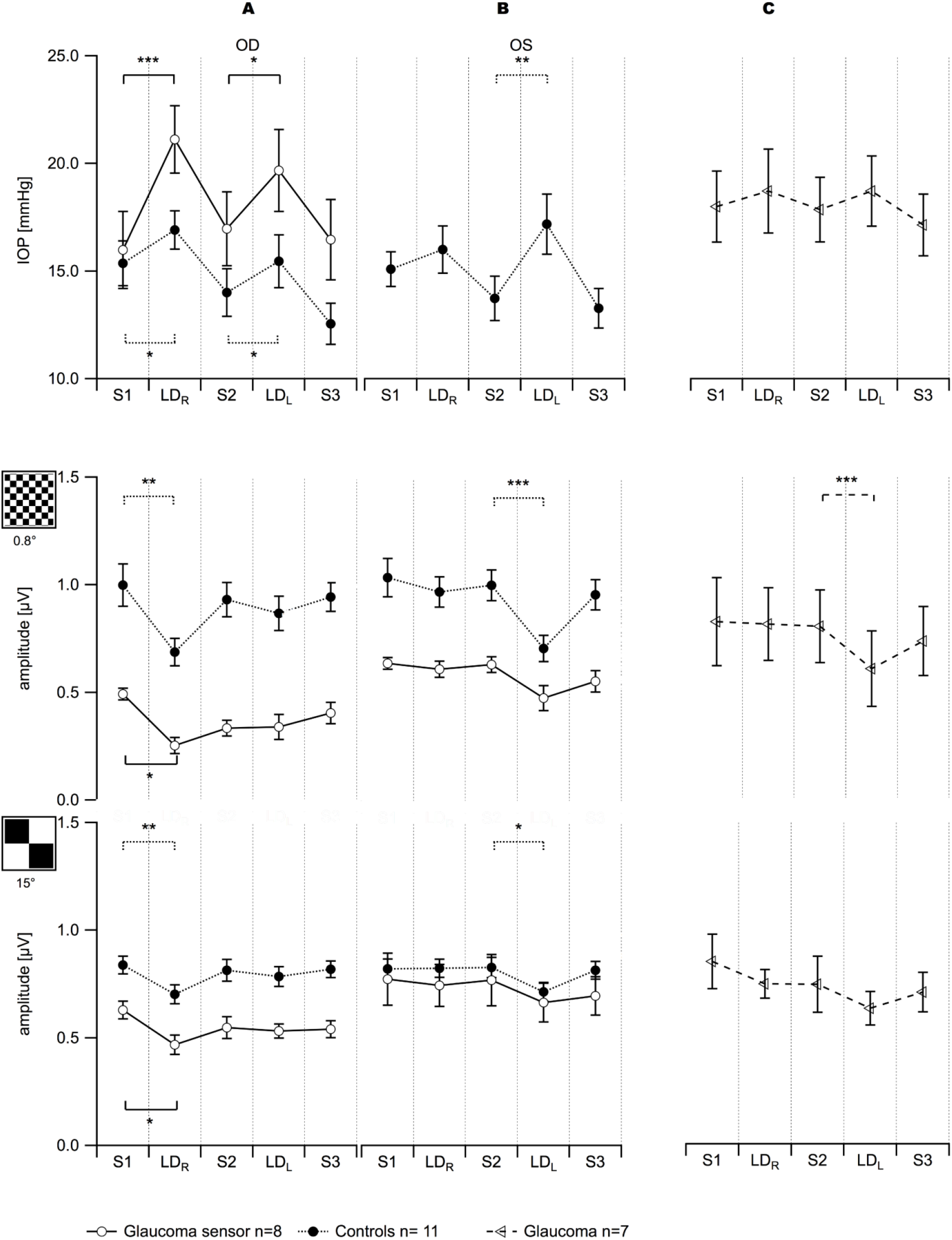
Mean±SEM IOP (top row) and Mean±SEM ssPERG data (0.8° checksize: middle row; 15° checksize: bottom row) for different postures (S1= sitting 1; LD_R_= right lateral decubitus; S2= sitting 2; LD_L_= left lateral decubitus; S3= sitting 3). T-test were performed for control right eyes (CON_RE_) vs GLA_IMP_-group’ right eyes and for control left eyes (CON_LE_) (A and B) vs left eyes GLA_LE_-group (C) and for conditions S1 vs LD_R_ and S2 vs LD_L_. * P value: < 0.05; ** P value < 0.01; *** P value < 0.001.

The above findings might be due either to the recorded right eye being the lower in LD_R_, alternatively it could be a sequential effect, as LD_R_ was the 2^nd^ and LD_L_ the 4^th^ condition. The former is clearly supported by Figure 4B, where the simultaneously recorded data from the left eye are depicted. Here, the IOP reductions were significant in controls for the lower eye during LDP. In CON_LE_, there was a significant increase of IOP during LD_L_ vs S2 by 3.5 mmHg (P < 0.01), while a trend for increased IOP for LD_R_ vs S1 did not reach significance (Figure 4B, Table 4). As the left eye of the GLA_IMP_-group was not glaucomatous for all subjects, we confirmed this effect in the GLA_LE_-group with glaucomatous left eyes (Figure 4 C and Table 3). In GLA_LE_, the statistical power of our design was not sufficient to resolve a significant difference of IOPs of left eye between sitting and LDP.

**Table 4.**
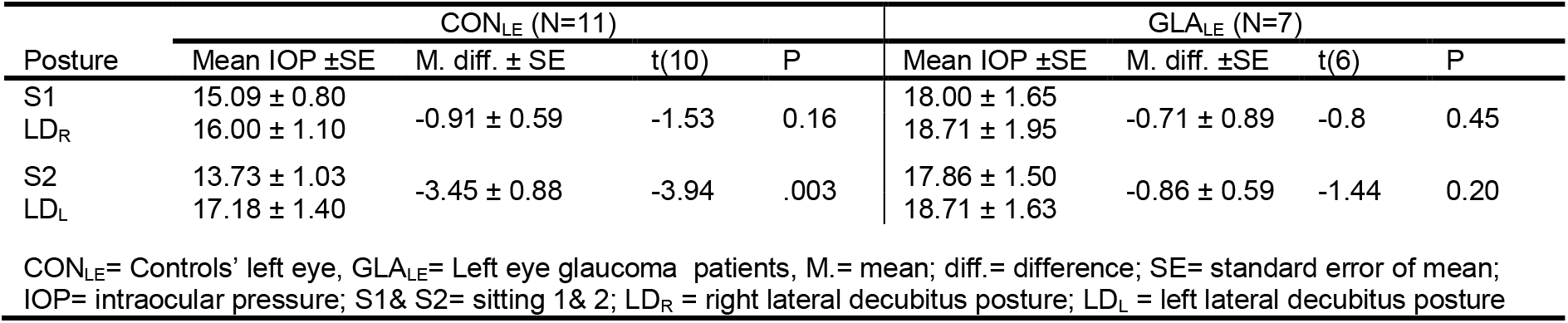
Intraocular pressure of left eye during different body postures

### Experiment 2 – Influence of postural change on PERG

For GLA_IMP_ right eye (Figure 4A) 0.8° and 15° PERG-amplitudes decreased [by 0.24 μV (P = 0.001) and 0.16 μV (P = 0.02), respectively] for LD_R_ vs S1, but not for LD_L_ vs S2. Similarly, for CON_RE_, 0.8° and 15° PERG amplitudes decreased for LD_R_ vs S1 by 0.31 μV (P = 0.001) and by 0.14 μV (P = 0.005), respectively, but not for LD_L_ vs S2 (Table 5). Similar effects, but as expected for LD_L_ vs S2, were evident for CON_LE_ as a significant amplitude decrease by 0.29 μV (P < 0.001) and 0.11 μV (P = 0.010) for 0.8° and 15° PERG-amplitudes, respectively, and for GLA_LE_ (Figure 4 C) as a significant amplitude decrease, which reached significance only for 0.8° (P < 0.001; Table 6).

**Table 5.** PERG amplitude changes of right eye during different body postures

| Check | Posture | CON<sub>RE</sub> (N=11) |  |  |  | GLA<sub>IMP</sub> (N=8) |  |  |  |
| --- | --- | --- | --- | --- | --- | --- | --- | --- | --- |
|  |  | M. Amplitude ± SE | M. diff. ± SE | t(10) | P | M. Amplitude ± SE | M. diff. ± SE | t(7) | P |
| 0.8° | S1 | 0.10 ± 0.10 |  |  |  | 0.49 ± 0.03 |  |  |  |
|  | LD<sub>R</sub> | 0.69 ± 0.06 | 0.31 ± 0.07 | 4.58 | 0.001 | 0.25 ± 0.04 | 0.24 ± 0.04 | 5.78 | 0.001 |
| 15° | S1 | 0.84 ± 0.04 |  |  |  | 0.63 ± 0.04 |  |  |  |
|  | LD<sub>R</sub> | 0.70 ± 0.04 | 0.14 ± 0.04 | 3.59 | 0.005 | 0.47 ± 0.05 | 0.16 ± 0.06 | 2.87 | 0.02 |
| 0.8° | S2 | 0.93 ± 0.08 |  |  |  | 0.33 ± 0.04 |  |  |  |
|  | LD<sub>L</sub> | 0.87 ± 0.08 | 0.06 ± 0.04 | 1.67 | 0.12 | 0.34 ± 0.06 | -0.01 ± 0.04 | -0.13 | 0.89 |
| 15° | S2 | 0.81 ± 0.05 |  |  |  | 0.55 ± 0.05 |  |  |  |
|  | LD<sub>L</sub> | 0.78 ± 0.05 | 0.03 ± 0.04 | 0.74 | 0.47 | 0.53 ± 0.03 | 0.02 ± 0.06 | 0.29 | 0.78 |

CON<sub>RE</sub>= Controls' right eye, GLA<sub>IMP</sub>= eyemate-IO sensor patients' right eye, M.= mean; diff.= difference; SE= standard error of mean; S1& S2= sitting 1& 2; LD<sub>R</sub> = right lateral decubitus posture; LD<sub>L</sub> = left lateral decubitus posture

**Table 6.** PERG amplitude changes of left eye during different body postures

| Check | Posture | CON<sub>LE</sub> (N=11) |  |  |  | GLA<sub>LE</sub> (N=7) |  |  |  |
| --- | --- | --- | --- | --- | --- | --- | --- | --- | --- |
|  |  | M. Amp.±<br>SE | M. diff. ±<br>SE | t(10) | P | M. Amp.±<br>SE | M. diff. ±<br>SE | t(6) | P |
| 0.8° | S1 | 1.04 ± 0.09 |  |  |  | 0.82 ± 0.20 |  |  |  |
|  | LD<sub>R</sub> | 0.97 ± 0.07 | 0.07 ± 0.05 | 1.318 | 0.217 | 0.81 ± 0.17 | 0.01 ± .09 | 0.12 | 0.91 |
| 15° | S1 | 0.82 ± 0.05 |  |  |  | 0.85 ± 0.13 |  |  |  |
|  | LD<sub>R</sub> | 0.82 ± 0.04 | -0.002 ± .04 | -0.065 | 0.949 | 0.75 ± 0.07 | 0.10 ± .07 | 1.48 | 0.19 |
| 0.8° | S2 | 1.01 ± 0.07 |  |  |  | 0.80 ± 0.17 |  |  |  |
|  | LD<sub>L</sub> | 0.71 ± 0.06 | 0.29 ± 0.06 | 5.167 | < .001 | 0.60 ± 0.18 | 0.20 ± .03 | 7.31 | < .001 |
| 15° | S2 | 0.83 ± 0.05 |  |  |  | 0.75 ± 0.13 |  |  |  |
|  | LD<sub>L</sub> | 0.71 ± 0.04 | 0.11 ± 0.04 | 3.180 | 0.010 | 0.63 ± 0.08 | 0.11 ± .09 | 1.31 | 0.24 |

CON<sub>LE</sub>= Controls' left eye, GLA<sub>LE</sub>= Left eye glaucoma patients, M.= mean; Amp.= amplitude; diff.= difference; SE= standard error of mean; S1& S2= sitting 1& 2; LD<sub>R</sub> = right lateral decubitus posture; LD<sub>L</sub> = left lateral decubitus posture

Taken together, LDP induced an IOP increase and PERG decrease predominantly in the lower eye.

## Discussion

In the present study we demonstrated that ssPERG recordings are feasible during simultaneous continous IOP measurements with the eyemate-IO sensor. This opens novel directions to investigate the relationship of IOP and ganglion-cell function. Our findings corroborate the hypothesis that LDP-induced IOP increase in the lower eye is accompanied by reduced ganglion response as determined with ssPERG, both in control and glaucoma eyes.

We observed in the LDP a significantly higher IOP increase in the lower eye compared to the reference condition, i.e. sitting, in both controls and GLA_IMP_. This is in accordance with previous studies, that demonstrated LDP induced IOP increase of the lower eye in comparison to either sitting and/or supine postures in either healthy subjects^10,11,40^ or untreated glaucoma patients^14,41^. The GLA_IMP_-group showed higher IOP fluctuations and rise than healthy subjects during postural change from sitting to LDP, which is in line with other studies that reported that higher LDP- or supine-induced IOP changes were associated with more progression or asymmetrical visual field defects of untreated and treated glaucoma patients^13,14,42–45^; however, others did not report a relationship of LDP-induced changes and glaucoma progression or laterality^15,46^. Our GLA_LE_-group showed no significant IOP change during LDP vs sitting. It should be noted, however, that in the present study the IOP measurements were performed with different devices in the GLA_LE_- and control group compared to the GLA_IMP_-group. Further, a study comparing the LDP-induced IOP changes in open and closed angle glaucoma and healthy controls did not report significant differences in IOP positional changes between the groups^47^. It is difficult to compare our findings to the aforementioned studies because of different methodologies used.

We observed a reversible reduction of ganglion cell responses as reflected by the ssPERG for LDP, especially in the lower eye. This is supported by a previous report on the effect of recumbent posture the ssPERG after 8 minutes of head-down body tilt (HDT) of −10° in a subpopulation of glaucoma suspects and early manifest glaucoma patients ^8,9^. It was also shown in a longitudinal study that glaucoma suspects at higher risk for glaucoma conversion had more susceptibility to moderate HDT changes ^9^. In contrast, to the authors’ knowledge, there is no prior study characterizing electrophysiological changes during LDP. As LDP does, in contast to supine posture, not require adapted stimulation set-ups, future studies using LDP might therefore open the possibility of a widespread use of the PERG-recordings during posture changes.

### Pathophysiology of IOP change in lateral decubitus

It was hypothesized that higher IOP in the lower eye during recumbent posture might be due to gravity and subsequent episcleral venous pressure changes^48^. In fact, biomechanical effects at the optic nerve head caused by choroidal vascular congestion and altered IOP and cerebrospinal fluid pressure gradient and increased IOP may contribute to PERG changes^49,50^. However, ocular perfusion pressure was not measured in the present study. It was also shown that artificial IOP elevation decreased the PERG signal in controls and ocular hypertension subjects ^51,52^.

### Practical considerations and potential applications

We provide proof-of-concept for telemetric IOP measurement during steady-state PERG recordings. The robustness of the PERG recordings is strongly related to the frequency-based amplitude measurements as this allows for the dissociation of the intrusions around 9.2 Hz from the stimulation related response at 15 Hz. As a matter of course, for a closer spacing of stimulation and readout frequencies (or their harmonics) than in the present study read-out intrusions might influence PERG measurements. As another consequence of the requirement of a frequency-related analysis to avoid reader intrusions, transient PERG recordings cannot be performed with simultaneous readout, unless other measures are taken, e.g. substantially longer averaging. Since we demonstrated LDP induced-IOP and -ganglion cell changes, it is of great promise to study continuous IOP variations during sleep position and in glaucoma patients and studying the feasibility of implementing LDP as a provocative test in glaucoma suspects in which those at risk of glaucoma conversions may show more postural induced IOP and ganglion cell responsiveness changes.

### Limitations and potential confounds

Potential confounds related to electrode placement, participant positioning and optics as well as other limitations were addressed. Artificial PERG changes induced by misalignment of the glasses or by electrode displacement during LDP were addressed by careful positioning. Checking the effect of astigmatism, i.e. the dependence of PERG amplitude changes on astigmatism, we did not find any significant correlations. Small sample size, due to the rarity of patients with an IOP-sensor implant, was another limiting factor in our study. Further, it should be noted the current observations may not reflect the real changes of IOP during sleep due to various physiological and environmental conditions that might additionally influence IOP.

### Conclusion

In this study, we demonstrate that continuous IOP monitoring with the eyemate-IO sensor can be used for simultaneous IOP measurements and ssPERG recordings. We report reduced ssPERG in the lower eye during lateral decubitus positioning and thus demonstrate the relation of IOP changes on retinal ganglion cell function. This opens the possibility to perform investigations to scrutinize the relationship of IOP and ganglion-cell function as tested in the present study. Further studies with bigger sample size are required to detail the observed effects and their relationship to glaucoma asymmetry and progression.

## Acknowledgements

This work was supported by European Union’s Horizon 2020 research and innovation programme under the Marie Sklodowska-Curie grant agreement (No. 675033) to Michael B. Hoffmann and Lars Choritz.

## Conflict of Interest Statement

None

